# Targeting HIV Env immunogens to B cell follicles in non-human primates through immune complex or protein nanoparticle formulations

**DOI:** 10.1101/2020.02.19.956482

**Authors:** Jacob T. Martin, Christopher A. Cottrell, Aleksandar Antanasijevic, Diane G. Carnathan, Benjamin J. Cossette, Chiamaka A. Enemuo, Etse H. Gebru, Yury Choe, Federico Viviano, Talar Tokatlian, Kimberly M. Cirelli, George Ueda, Jeffrey Copps, Torben Schiffner, Sergey Menis, William R. Schief, Shane Crotty, Neil P. King, David Baker, Guido Silvestri, Andrew B. Ward, Darrell J. Irvine

**Author notes:** Equal contributions (Jacob T. Martin, Christopher A. Cottrell, Aleksandar Antanasijevic).

## Abstract

Following immunization, high affinity antibody responses develop within germinal centers (GCs), specialized sites within follicles of the lymph node (LN) where B cells proliferate and undergo somatic hypermutation. Antigen availability within GCs is important, as B cells must acquire and present antigen to follicular helper T cells to drive this process. However, recombinant protein immunogens such as soluble HIV envelope (Env) trimers do not efficiently accumulate in follicles following traditional immunization. Here we demonstrate two strategies to concentrate HIV Env immunogens in follicles, via the formation of immune complexes (ICs) or by employing self-assembling protein nanoparticles for multivalent display of Env antigens. Using rhesus macaques, we show that within a few days following immunization, free trimers were present in a diffuse pattern in draining LNs, while trimer ICs and Env nanoparticles accumulated in B cell follicles. Whole LN imaging strikingly revealed that ICs and trimer nanoparticles concentrated in as many as 500 follicles in a single lymph node within 2 days after immunization. Imaging of LNs collected 7 days post-immunization showed that Env nanoparticles persisted on follicular dendritic cells in the light zone of nascent germinal centers. These findings suggest that the form of antigen administered in vaccination can dramatically impact localization in lymphoid tissues and provides a new rationale for the enhanced immune responses observed following immunization with immune complexes or nanoparticles.

## Background

The development of immunogens/immunization regimens capable of eliciting broadly neutralizing antibodies (bNAbs) that can recognize diverse viral strains is thought to be an important approach to achieve an effective HIV vaccine ^1,2^. The sole target for neutralizing antibodies on the virus surface is the Env protein, a homotrimer consisting of three copies of gp120 and gp41 subunits that are associated by non-covalent interactions. The development of methods to produce stable, recombinant, trimer immunogens based on the ectodomain of Env have catalyzed HIV vaccine efforts ^3–10^. However, optimizing immunization strategies to promote high-affinity antibody responses against trimer immunogens remains an important goal. B cell affinity maturation occurs in germinal centers formed within B cell follicles in lymph nodes, and the availability of antigen to B cells within GCs plays a critical role in regulating the outcome of immunization ^11–13^. Hence, the efficient trafficking of Env trimer immunogens to follicles following immunization is essential to promote optimal humoral responses to HIV.

Upon immunization, soluble immunogens are rapidly bound by antibodies present in the tissue, forming immune complexes (ICs) that can subsequently traffic into lymphatic vessels and downstream draining lymph nodes (LNs). In antigen-naïve animals, IC formation can be mediated by natural pentameric IgM (nIgM) that binds antigen with low affinity but moderate to high avidity ^14,15^. Antigen-antibody ICs are subsequently opsonized by complement, ultimately resulting in the covalent attachment of the complement protein C3d ^14,15^. As the IC-C3d complexes enter the draining LN, they are captured by subcapsular sinus (SCS) macrophages via complement receptors and are transported across the subcapsular sinus floor into the B cell follicle ^16–18^. Non-antigen specific, naïve B cells capture IC-C3d complexes from the basolateral surface of SCS macrophages using complement receptor 2 (CR2) and deposit them onto the surface of follicular dendritic cells (FDCs), where they are subsequently available to participate in the GC reaction ^17^. FDCs serve as antigen depots that are capable of storing and displaying antigen for months ^19^.

Soluble HIV Env immunogens do not follow the above canonical pathway for entering the B cell follicle ^20^. Most likely due to the high level of glycosylation and the low capacity to form avidity-enhanced interactions, soluble Env trimers do not efficiently bind nIgM or activate complement ^21^. Instead, they enter the LN as free antigens and are captured by interfollicular channel (IFC) macrophages via cell surface SIGN-R1 receptors ^20^. Antigen-specific naïve B cells have an opportunity to interact with captured antigen on the surface of IFC macrophages as the B cells exit the circulatory system on their way to the B cell follicle. Additionally, IFC macrophages extend antigen-bearing cellular processes into the B cell follicle where antigen-specific follicular B cells can capture antigen along the edge of the follicle ^20^. This alternative antigen trafficking pathway allows for the initial activation of B cells but does not provide additional antigen needed for repeated cycles of the GC reaction ^20^. Presumably, the initial activation of naïve B cells results in a production of short-lived plasmablasts, which secrete antibodies capable of forming ICs with soluble Env immunogens that can eventually be deposited on FDCs ^22^. This delay in Env immunogen trafficking to FDCs could potentially result in a loss of antigenic integrity, due to degradation of the immunogen by endogenous proteases, promoting dominance of antibody responses focused on non-neutralizing epitopes ^21,23^.

These issues motivate the exploration of methods to promote rapid delivery of Env immunogens to follicles, such as alternative approaches to engage the natural complement-mediated pathway for delivery of ICs to follicles. Fusion of antigens with multiple copies of recombinant C3 has been shown to enhance humoral immunity ^24^, but this strategy has not been widely applied and may be difficult to implement generally. Another approach is to form synthetic immune complexes with recombinant monoclonal antibodies ^25–28^. Immunization with ICs in animal models has been shown in some cases to enhance IgG titers and duration, and to affect which epitopes are targeted ^27–30^. Early-stage clinical trials of IC immunization have shown this approach to also be safe ^31,32^. Passive administration of broadly neutralizing antibodies (bnAbs) against HIV Env is now being investigated in numerous clinical trials with good safety profiles reported for a variety of bnAbs ^33–35^, suggesting a pathway for rapid clinical translation of IC immunization against HIV.

Recently, we discovered another approach for promoting complement decoration of immunogens that is independent of the classical pathway of complement activation by antibodies. Immunization with gp120 monomer or gp140 trimer antigens fused to proteins that form nanoparticles through self-assembly led to rapid concentration of these immunogens on LN FDCs as early as 24 hr after injection ^36^. Delivery of these particulate antigens to follicles was dependent on C3 and complement receptors. Complement activation was mediated by mannose-binding lectin (MBL), which bound with high avidity to the densely glycosylated HIV Env nanoparticles but not the soluble forms of these immunogens. MBL binding enhanced GC responses, increased the production of long-lived plasma cells in bone marrow, and increased serum antibody responses ^36^.

Both IC and nanoparticle immunization have been reported to enhance the immunogenicity of subunit vaccines in small and large animals, but the mechanisms underlying these enhancements remain poorly understood – especially in large animals such as rhesus macaques that are similar to humans. Notably, these two approaches enhance complement binding to the immunogen, which may alter antigen fate in LNs. Here, we investigated the impact of IC formation and nanoparticle presentation on HIV Env immunogen trafficking in lymphoid tissues of non-human primates. First, to enhance the formation of ICs, we passively administered, prior to immunization, a high affinity macaque-derived monoclonal antibody that targets the base of the soluble HIV Env trimer immunogen BG505 SOSIP. Second, to enhance mannose-binding lectin (MBL) binding and subsequent complement deposition, we multimerized the BG505 SOSIP HIV Env trimer immunogen by genetically fusing it to one of the components of a self-assembling, two-component tetrahedral nanoparticle ^37,38^. We applied these two immunization strategies in rhesus macaques, evaluating their impact on immunogen accumulation in draining LNs and localization within LN tissues. We find that both approaches promote the concentration of Env immunogens in B cell areas, with antigen accumulation in hundreds of follicles within a single lymph node. Nanoparticle immunogens were further shown to persist in follicles for at least a week, with a distribution suggesting accumulation on FDCs making up the light zone of nascent geminal centers. Thus, IC and NP immunizations can dramatically alter the localization of antigen in lymph nodes, providing a rationale for their application in vaccine design platforms to promote humoral immunity against HIV and other infectious diseases.

## Results

### Characterization of a rhesus macaque monoclonal antibody which recognizes the base of BG505 SOSIP Env trimers

The mAb RM19R was isolated from a BG505 SOSIP trimer-immunized rhesus macaque and shown to bind to the base of the trimer by single particle negative stain electron microscopy (NS-EM) ^39^. Fragments antigen binding (FAbs) of RM19R bind to the BG505 SOSIP trimer with an affinity of 0.55 nM, as determined by biolayer interferometry (BLI) (Fig. 1A). RM19R IgG also forms immune complexes with the BG505 SOSIP *in vitro* as shown in Fig. 1B. To determine the molecular details of the RM19R epitope, we solved a 3.7 Å resolution cryo-EM structure of BG505 SOSIP in complex with RM19R Fabs (Fig. 1C and **Supplementary Fig. 1 A-D**). RM19R recognizes a quaternary epitope spanning two gp41 protomers and a single gp120 protomer that has 1316 Å² of buried surface area at the interface (Fig. 1C-D). RM19R uses two tyrosine residues at the tip of its CDR H3 to wedge between the C-terminal region of gp120 and the C-terminus of gp41 in the adjacent protomer (Fig. 1D). Binding of RM19R causes the last 7 residues (Q658 to D664) of the gp41 HR2 helix to unwind (Fig. 1D), but despite this conformational change, the complex is highly stable at room temperature for more than 24 hours (**Supplementary Fig. 1E**). The light chain of RM19R also makes contact with the C5 region of gp120, and an arginine residue at position 500 in gp120 inserts between the heavy and light chains of RM19R (Fig. 1D). Consistent with the observations in the cryo-EM structure, binding of RM19R is eliminated by combining the mutations R500A and Q658K (Fig. 1E).

**Figure 1.**
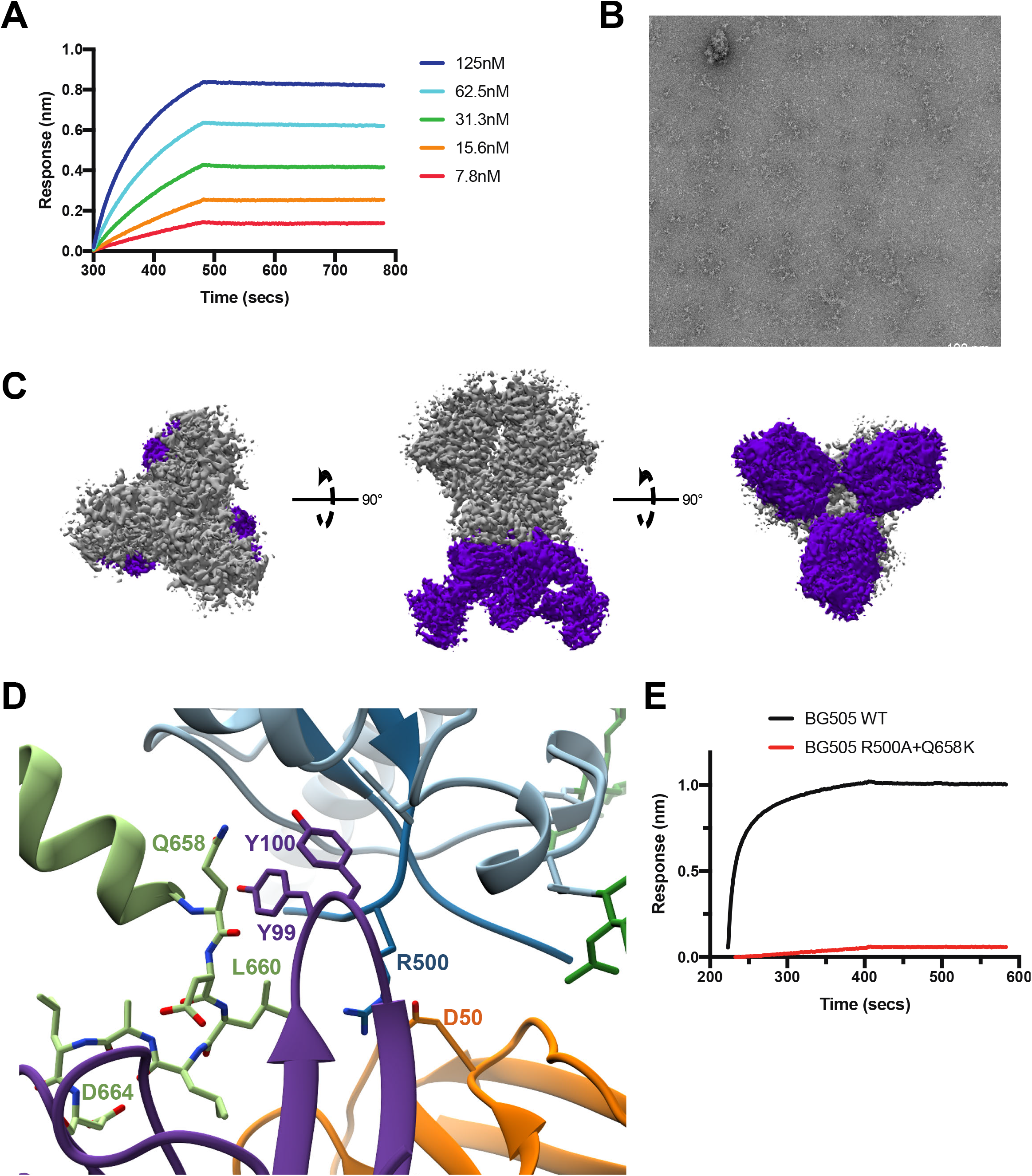
RM19R mAb binds to the base of Env trimer. (A) BLI curves for immobilized RM19R Fab binding to BG505 SOSIP.v5.2 trimer as a function of SOSIP concentration. (B) Negative stain EM image of BG505 SOSIP.v5.2 trimer/RM19R IgG in vitro immune complexes. (C) CryoEM 3D reconstruction (3.7 Ǻ resolution) of the BG505 SOSIP.v5.2.N241.N289 trimer/RM19R Fab complex. BG505 trimer density in gray with the RM19R Fabs density in purple. (D) Binding pocket of RM19R CDR H3 with BG505, showing interactions between R500 and Q658 of BG505 with Y99 and Y100 of RM19R RM19R heavy chain (purple), RM19R light chain (orange), gp120 (blue), gp41 (light blue), glycans (green), and the adjacent gp41 (light green). (E) BLI curves of immobilized RM19R IgG to BG505.v5.2 (BG505 WT, black) or BG505 SOSIP.v5.2.N241.N289.R500A.Q658K (BG505 R500A+Q658K, red). Binding to RM19R is abolished by the combined mutations R500A and Q658K.

### Immune complex formation redirects soluble Env immunogens to B cell follicles in LNs of mice and non-human primates

To determine the impact of immune complex formation between BG505 SOSIP trimers and the base-directed RM19R antibody *in vivo*, we first analyzed trafficking of labeled RM19R and trimer by whole-tissue fluorescence imaging in mice. Groups of animals were passively immunized intravenously (i.v.) with doses of RM19R antibody ranging from ~0.25 mg/kg to 10 mg/kg, or PBS as a control. Twenty-four hours later, the animals were subcutaneously (s.c.) administered fluorescently-labeled BG505 SOSIP trimer. Twenty-four hours following immunization, the animals were sacrificed and draining inguinal LNs were removed and fixed for imaging. Whole-tissue fluorescence imaging using a Typhoon flatbed fluorescence scanner showed peak trimer accumulation in draining LNs at 1 day after immunization (Fig. 2A). Confocal microscopy imaging of whole lymph nodes after tissue clearing and staining for FDCs using an anti-CD35 antibody revealed substantial differences in the localization of the trimer immunogen in the presence of RM19R. BG505 trimer colocalized with B cell follicles in the presence of the RM19R in a dose-dependent manner (Fig. 2B). This observation suggested that higher concentrations of antigen-specific antibodies resulted in more complete incorporation of the soluble antigen into immune complexes, which accumulated on FDCs.

**Figure 2.**
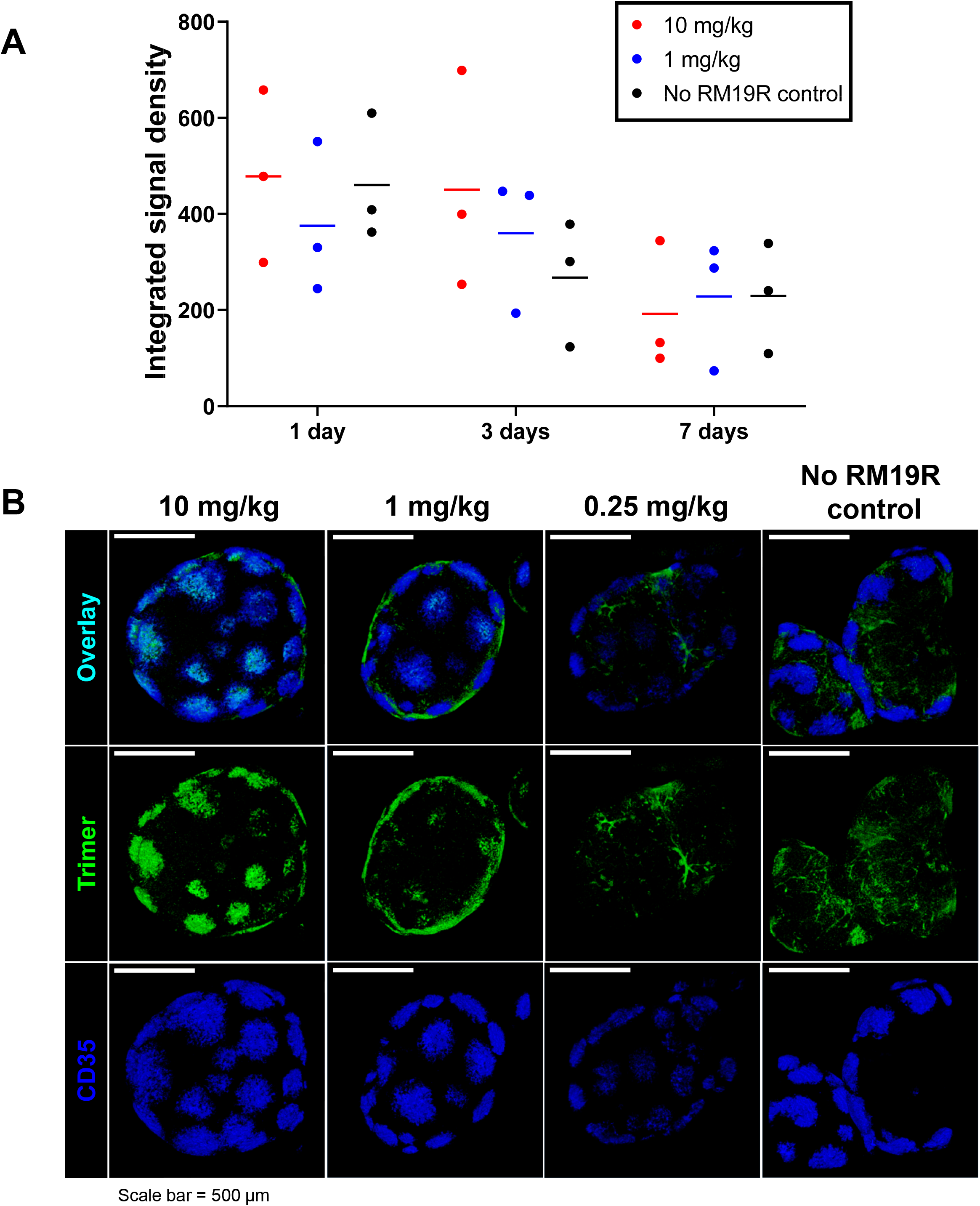
Immunization of mice with Env trimer in the presence of passively transferred antibody leads to rapid antigen accumulation in follicles. Balb/c mice (n = 3/group) were injected i.v. with different doses of RM19R mAb followed 24 hr later by s.c. immunization with 5 µg AlexaFluor568-labeled BG505 trimer. Draining inguinal LNs were collected at various times after immunization for whole-tissue imaging. (A) Total trimer fluorescent signal in whole fixed LNs at 1 to 7 days post immunization, as measured by flatbed fluorescence scanner. (B) 3D projections of whole cleared inguinal LNs 24 hr after immunization, imaged by confocal microscopy to a depth of 300 µm. Scale bars 500 µm.

We next carried out analogous experiments in rhesus macaques to compare antigen trafficking in large animals to our observations in mice. For this study we labeled both the RM19R mAb and the BG505 SOSIP trimer, to enable tracking of both antigen and antibody. Pairs of non-human primates (NHPs) were injected i.v. with fluorescently labeled RM19R mAb at 2.0 mg/kg or 0.2 mg/kg, and 24 h later, all animals received s.c. injections of 50 µg BG505 SOSIP immunogen in each thigh; a control group was immunized s.c. with the trimer in the absence of RM19R passive transfer. Three days after antigen injection, animals were sacrificed and inguinal, iliac, axillary, popliteal, and mesenteric lymph nodes were collected for analysis. Total RM19R and trimer accumulation in LNs were first measured by IVIS fluorescence imaging of excised tissues. To account for variability between animals in the number of LNs at a given site, we summed fluorescence signals for LNs collected at a common drainage location (e.g., all iliac LNs, or all axillary LNs). As expected from prior studies of antigen trafficking from this injection site in NHPs ^3^, inguinal and iliac LNs were the primary draining LNs where SOSIP trimer accumulated (Fig. 3A). In line with the observations in mice, the total amount of trimer immunogen accumulating in these LNs was unaffected by the presence of passively transferred RM19R (Fig. 3A). In contrast, RM19R antibody showed preferential accumulation only in antigen-draining LNs, in an antibody dose-dependent manner (Fig. 3B).

**Figure 3.**
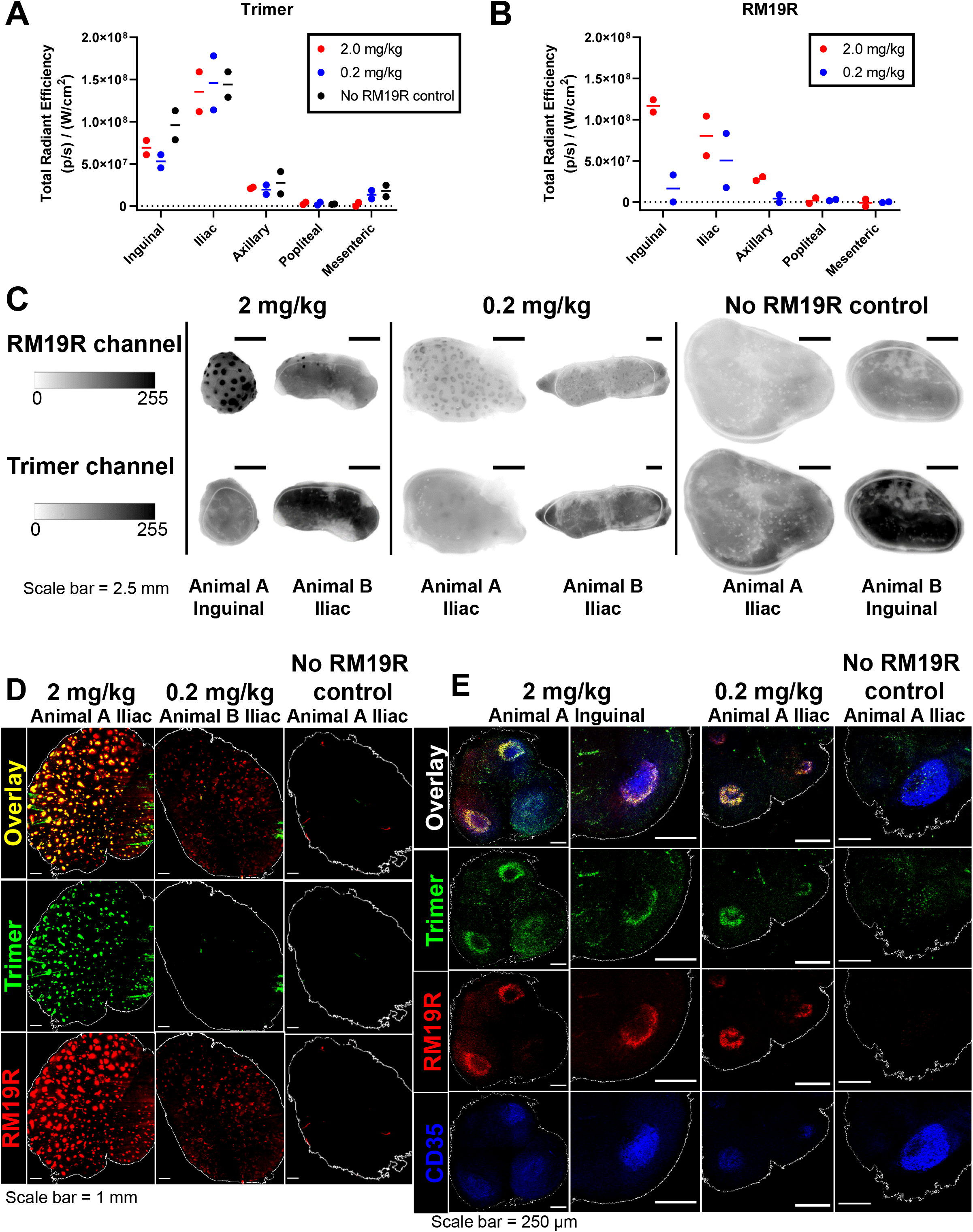
Immunization of NHPs with Env trimer in the presence of passively transferred antibody leads to rapid antigen accumulation in follicles. Rhesus macaques (n = 2/group) were injected i.v. with different doses of AlexaFluor647-labeled RM19R mAb followed 24 hr later by s.c. immunization with 50 µg AlexaFluor568-labeled BG505 trimer in each mid thigh. LNs from indicated locations were harvested 3 days after immunization. (A) Total fluorescent signal corresponding to trimer from all LNs from each animal at a particular site, as measured by IVIS. (B) Total fluorescent signal corresponding to RM19R antibody from all LNs from each animal at a particular site, as measured by IVIS. (C) Selected images of whole uncleared LNs from fluorescent flatbed scanner imaging in each channel. Scale bar represents 2.5 mm. (D) Maximum projections of whole cleared LNs from draining LN locations for each group, as imaged by light sheet microscopy. Scale bar represents 1 mm. (E) Single planes of uncleared 100 µm-thick LN sections immunofluorescently-stained for FDCs, as imaged by confocal microscopy. Scale bar represents 250 µm.

Fluorescence imaging of fixed whole LNs at low magnification using a Typhoon flatbed fluorescence scanner showed an inhomogeneous distribution of trimer and RM19R mAb within the tissues (Fig. 3C). Antigen-draining LNs from animals receiving RM19R prior to trimer injection showed focal accumulations of the mAb across the tissue that we hypothesized represented B cell follicles; this focal accumulation appeared to be more intense in animals receiving the high dose of mAb. Inhomogeneity in the distribution was also detectable for the BG505 SOSIP trimer antigen in some lymph nodes from animals that received the 2 mg/kg dose of the RM19R antibody (Fig. 3C, 2 mg/kg animal A inguinal LN). To obtain further insight into the distribution of antigen within the tissues, iliac LNs from each group were processed for whole-organ tissue clearing ^40,41^ and then imaged by light sheet microscopy. As shown in Fig. 3D and **Supplementary Video 1**, imaging of the cleared organs revealed strong colocalization of trimer and RM19R for animals receiving 2 mg/kg of the antibody, in as many as 500 follicles within a single LN. In accordance with the scanner images, the low dose group exhibited RM19R antibody accumulation in follicles, but colocalization of the trimer signal was less pronounced. No fluorescent signal above background was observed in the cleared samples from animals that did not receive passive transfer of the RM19R antibody. The fact that these control tissues showed substantial trimer signal in IVIS/Typhoon measurements prior to clearing suggests that the tissue clearing process extracted non-immobilized trimer immunogen from the tissues. In contrast, in the presence of the antibody, immune complex formation led to accumulation of ICs in focal areas which prevented removal of the trimer by the solvent washes used in tissue clearing.

To investigate whether the apparent focal accumulation of ICs was in fact due to deposition on the FDC network of individual B cell follicles, we employed immunofluorescent staining of 100 µm tissue sections. Confocal microscopy of stained sections of the LNs revealed bands of antigen and RM19R antibody signal around the perimeters of CD35^+^ FDC networks in multiple locations throughout each LN, corresponding to multiple B cell follicles (Fig. 3E). These ring-like accumulations of mAb and trimer correspond to the cross-sectional views through the cup-like, fluorescent 3D structures observed in the light sheet imaging of whole tissues (**Supplementary Video 2**). Follicle accumulation of mAb/trimer was most apparent in animals that received the highest dose of RM19R, but was consistently observed in the low-dose samples as well. These tissue samples were not subjected to clarification treatment prior to imaging, and antigen fluorescence was still detected in the samples from animals immunized with trimer in the absence of RM19R mAb. However, the pattern of antigen distribution was much more diffuse, and not concentrated on the FDC network (Fig. 3E). Thus, immunization with BG505 SOSIP trimer antigens in the presence of the trimer base-specific antibody RM19R greatly increased antigen accumulation in follicles.

### Production and characterization of a modular Env trimer protein nanoparticle

We next sought to compare the lymph node localization of ICs with a second strategy for promoting follicular localization, formulating Env trimers as protein nanoparticles ^36^. Two-component, self-assembling nanoparticles represent a promising platform for presentation of HIV Env immunogens ^37,38,42,43^. In this platform, the full length ectodomain of Env (gp140) is genetically fused to the trimeric component of the nanoparticle and the fusion (antigen-bearing component) can be expressed and purified independently using an equivalent set of approaches applied with free soluble ectodomains. The antigen-bearing component is then combined with the corresponding second building block (assembly component) to form a nanoparticle of well-defined geometry (Fig. 4A). This system offers more flexibility and control compared to alternative approaches based on natural protein scaffolds (e.g., ferritin, E2p, lumazine synthase) or nanoparticles prepared from synthetic polymers or inorganic materials.

**Figure 4.**
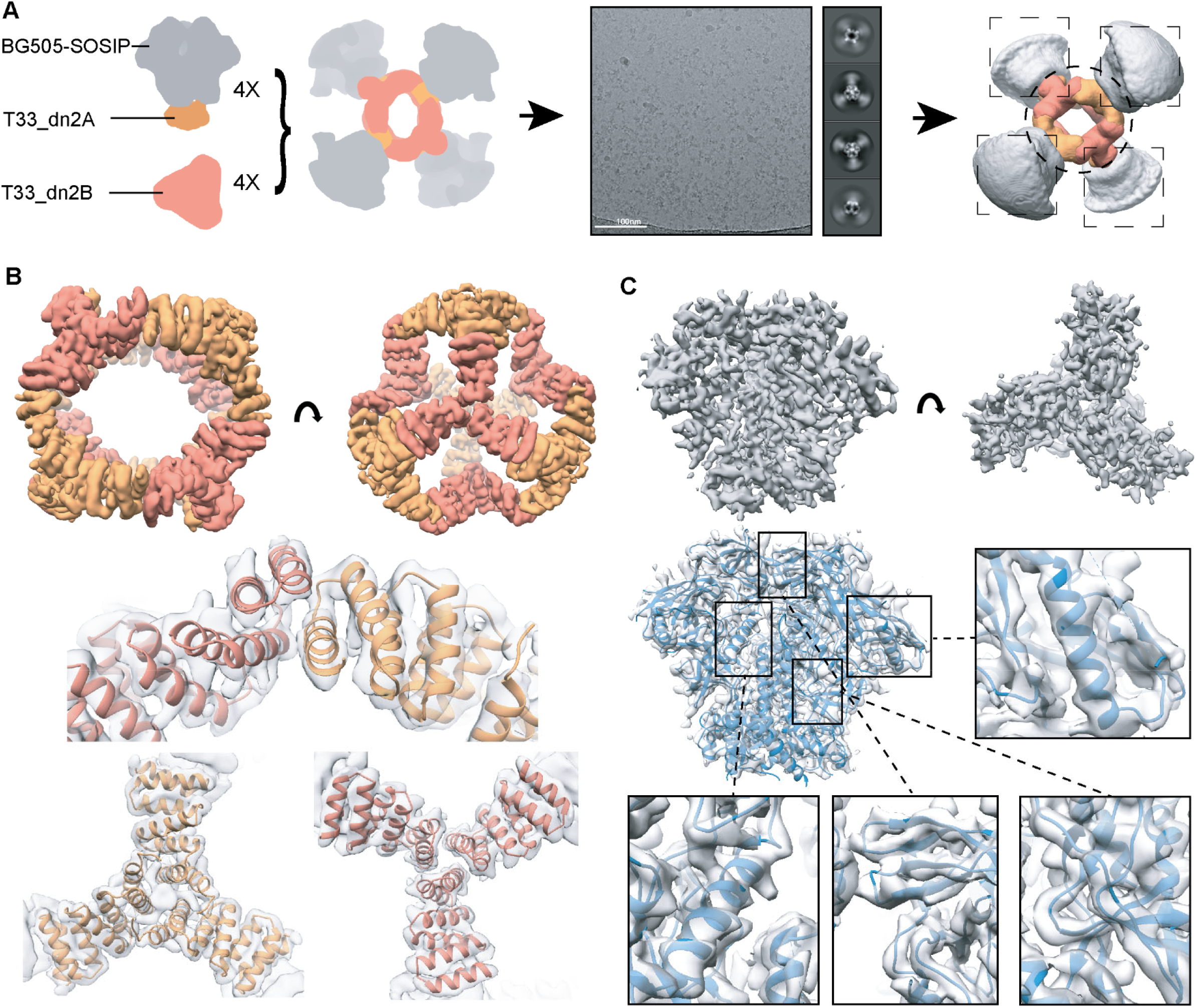
BG505 SOSIP-T33_dn2 nanoparticle self assembles in a tetrahedral arrangement. (A) Cryo-EM analysis of the BG505 SOSIP-presenting T33_dn2 nanoparticle. (B) Post-processed cryo-EM map of the nanoparticle core and the fit of refined model to the reconstructed density (T33-DN2-A and T33-DN2-B components are colored in orange and salmon, respectively). (C) Post-processed cryo-EM map of the nanoparticle-presented BG505 SOSIP trimer following subparticle analysis and model to map fit of the refined model.

We employed a tetrahedral nanoparticle, T33_dn2 ^37,38^, which consists of two trimeric building blocks (four copies of each), and can present four SOSIP trimers. A variant of BG505 SOSIP, engineered to have increased soluble expression (see methods for details), was fused to component A of the nanoparticle (T33_dn2A). For particle production, BG505 SOSIP-T33_dn2A was combined with equimolar amounts of the assembly component (T33_dn2B) and incubated for 24 hours, leading to >90% assembly efficiency (**Supplementary Fig. 2A**). SDS-PAGE analysis confirmed the presence of both BG505 SOSIP-T33_dn2A and T33_dn2B components in the purified nanoparticle sample (**Supplementary Fig. 2A**). Negative stain EM analysis revealed that the purified nanoparticles were highly homogeneous (**Supplementary Fig. 2B**).

BG505 SOSIP-T33_dn2 nanoparticles were then subjected to cryo-electron microscopy for structural analysis (Fig. 4, **Supplementary Fig. 3** and **Supplementary Table I**). An initial cryo-EM reconstruction of the entire nanoparticle verified the appropriate assembly to the target tetrahedral architecture with four BG505 SOSIP trimers flexibly linked to the nanoparticle core (Fig. 4A). The particle diameter is ~37 nm and the average apex-apex distance between the BG505 SOSIP trimers is 31 nm. Due to the flexible nature of the linker connecting the BG505 SOSIP antigen and T33_dn2A scaffold, high resolution reconstruction of the entire nanoparticle as a single model could not be performed. However, focused refinement in combination with partial signal subtraction allowed us to treat the nanoparticle core (consisting of T33_dn2A and B) and the displayed BG505 SOSIP trimers as separate subparticles and analyze them independently. Using the approach described in the methods section and **Supplementary Fig. 3A** we reconstructed a 4.6 Å resolution map of the nanoparticle core (Fig. 4B) and a 4.5 Å resolution map of the BG505 SOSIP trimer (Fig. 4C, **Supplementary Fig. 3A**). A combination of Rosetta relaxed refinement and manual refinement in Coot was applied to generate the atomic models (**Supplementary Table II**, Fig. 4B-C). The refined model of the T33_dn2 nanoparticle core revealed a high degree of correlation with the design model predicted by Rosetta at the backbone level. When compared to Rosetta models, Cα RMSD values for the trimeric T33_dn2A and T33_dn2B components were 1.20 Å and 0.81 Å, respectively (**Supplementary Fig. 3B**). The C-terminal helix in T33_dn2B was not resolved in the EM map, suggesting a high degree of flexibility which may be at least partially influenced by the 6xHis-tag immediately following this sequence. Importantly, the structure of the scaffolded BG505 SOSIP revealed that the antigen remains in the native-like, pre-fusion state and that nanoparticle incorporation did not interfere with its structure. Extensive antigenicity analysis of the BG505 SOSIP-T33_dn2 nanoparticle reported elsewhere ^37,38^ also supports this conclusion.

### Mannose-binding lectin recognizes BG505 Env trimer nanoparticles

We previously showed that Env trimers displayed on a ferritin nanoparticle core could be recognized by the innate immunity protein mannose-binding lectin (MBL), via the dense glycan coat on the trimer surfaces ^36^. We therefore evaluated the capacity of free trimeric BG505 SOSIP or BG505 SOSIP presented on tetrahedral T33_dn2 nanoparticles to interact with mannose-binding lectin *in vitro*. Recombinant human MBL-2 was incubated with BG505 SOSIP or BG505 SOSIP-T33_dn2 in buffer containing 2 mM CaCl_2_ and imaged using NS-EM as described in the methods section. NS-EM analysis confirmed the presence of high-molecular weight species in both samples (**Supplementary Fig. 4A**). However, EM data interpretation is complicated by the fact that MBL-2 itself exists in different oligomeric forms, some of which are ~20 Å in diameter or larger (**Supplementary Fig. 4A**). MBL-2 and antigen complex solutions were therefore also analyzed by size exclusion chromatography (SEC) (**Supplementary Fig. 4B, C**). Env trimer alone eluted primarily as a single peak in SEC (**Supplementary Fig. 4B**). When mixed with MBL-2, a small peak at low elution volume was detected, indicative of a limited formation of MBL/trimer aggregates. Surprisingly, MBL-2 also appeared to induce partial trimer disassembly and an increase in monomer/dimer species (**Supplementary Fig. 4B**). By contrast, BG505 SOSIP-T33_dn2 nanoparticles appeared to co-assemble into a greater proportion of higher molecular weight species when incubated with MBL-2 (**Supplementary Fig. 4C**). The aggregate fractions were subjected to NS-EM and SDS-PAGE analysis (**Supplementary Fig. 4D-E**). NS-EM analysis revealed the existence of large complexes of different sizes with discernable nanoparticles that appear to be clustered. SDS-PAGE confirmed the presence of both nanoparticle and MBL-2 in the aggregate sample. These biophysical measurements suggest that despite the modest antigen valency of the BG505 SOSIP-T33_dn2 nanoparticle, it can readily interact and co-aggregate with MBL that will be present in interstitial fluid.

### Nanoparticle trimer immunogens accumulate in follicles of NHPs following immunization

We next designed an experiment to compare antigen trafficking of soluble BG505 SOSIP MD39 trimer ^9^ and BG505 SOSIP-T33_dn2 nanoparticle immunogen in rhesus macaques. The immunogen dose was normalized based on the total amount of BG505 SOSIP (100 µg and 142 µg per dose per animal for free trimer and nanoparticle, respectively). Each animal received half of the total dose of AF647-labeled immunogen mixed with an ISCOMs-like saponin adjuvant s.c. in each of their left and right inner thighs. Groups of three animals were sacrificed either two or seven days post-immunization (Fig. 5A), and LNs from primary draining sites (inguinal and iliac) as well as distal sites (axillary), were harvested for imaging analysis.

**Figure 5.**
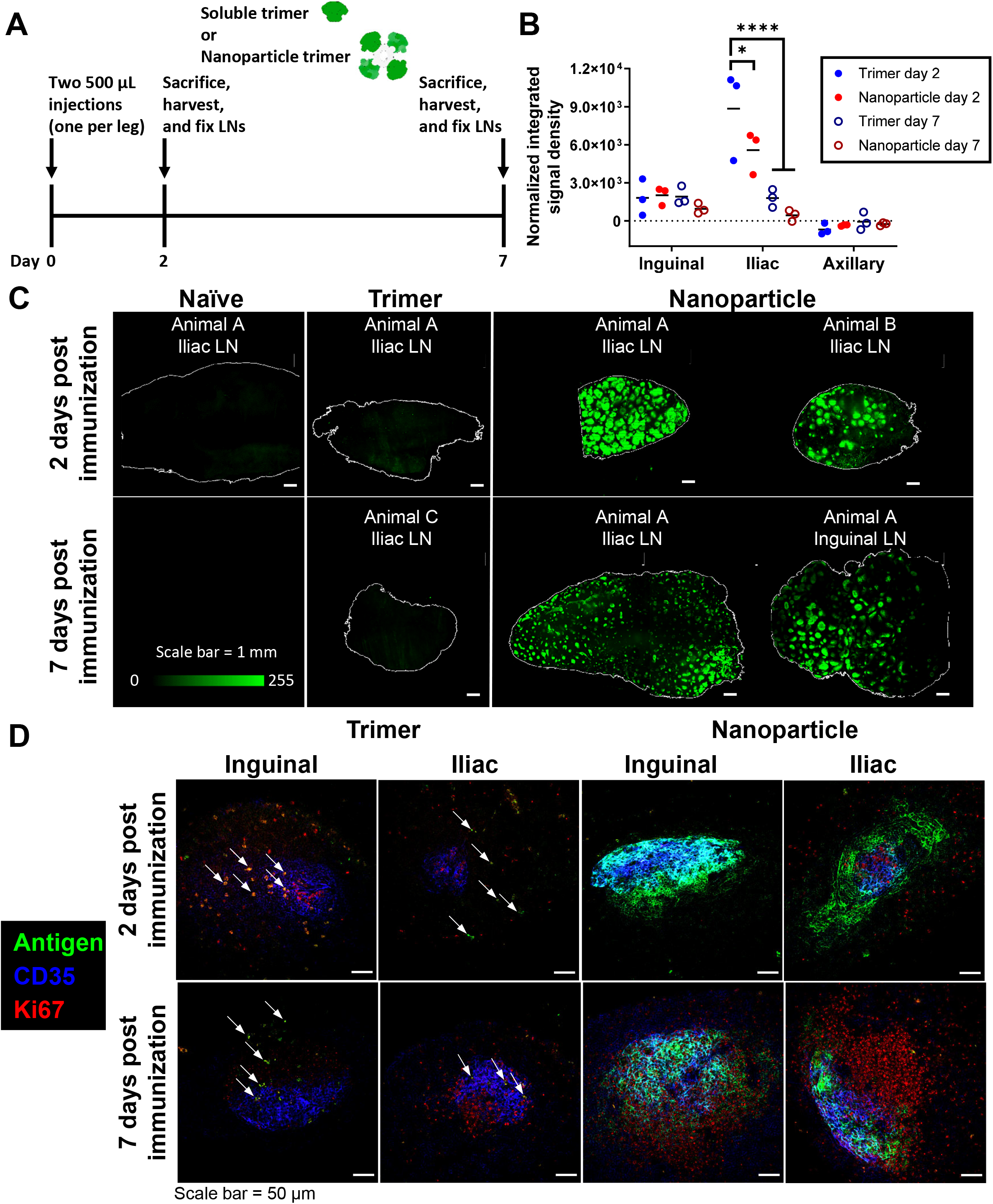
Immunization of NHPs with nanoparticle leads to rapid antigen accumulation in follicles without passively transferred antibody. (A) Schematic of the timeline of the study. (B) Total fluorescent signal corresponding to trimer or nanoparticle immunogen, normalized by degree of labeling per trimer. Signal is the summation from all LNs from a particular site for each animal, as measured by fluorescent flatbed scanner. * p = 0.0245, **** p = <0.0001 by two-way ANOVA. (C) 3D projections of whole cleared LNs from draining LN locations for each group, as imaged by light sheet microscopy. Scale bar 1 mm. (D) Single planes of uncleared 100 µm-thick LN sections immunofluorescently-stained for FDCs and Ki67, as imaged by confocal microscopy. Scale bar 50 µm.

We first analyzed the total antigen content of fixed lymph nodes using the Typhoon scanner, summing the total antigen fluorescence signal of all lymph nodes at each collection site as performed in the immune complex experiments. These measurements revealed a 37% lower accumulation of the antigen in the proximal iliac draining LNs for animals immunized with the nanoparticle compared to the soluble trimer immunogen at two days post immunization (Fig. 5B). We next selected individual antigen-positive LNs from each group (or naïve LN controls) and cleared them for light sheet imaging. Similar to results from the IC experiment, we observed little to no fluorescence above background in cleared organs from animals receiving soluble BG505 SOSIP trimer and adjuvant at either two or seven days post immunization (Fig. 5C and **Supplementary Video 3**). In striking contrast, all LNs from animals that received BG505 SOSIP presented on the T33_dn2 nanoparticle displayed bright and obvious focal accumulations of the immunogen throughout the lymph nodes, with up to 500 such follicular deposits observed in a single LN (Fig. 5C and **Supplementary Video 3**).

Next, histological sections of the main draining LNs were stained with anti-CD35 to identify the follicular dendritic cell networks, to confirm that the focal accumulations observed in the LNs of immunized animals are in fact in B cell follicles. We observed the accumulation of nanoparticle immunogen, but not free trimer, in and around B cell follicles (Fig. 5D). Free trimer was instead detected at low levels in the interior medullary regions of LNs, which may indicate uptake by DC-SIGN^+^ macrophages, as suggested by Park et al. ^20^. By co-staining the histological sections of draining lymph nodes with the proliferation marker Ki67, we detected germinal center induction in both trimer and nanoparticle-immunized animals at day 7 (Fig. 5D). However, free trimer was only detected in follicles very infrequently, suggesting very low levels of antigen in the GCs, while substantial levels of the nanoparticle-presented trimer were found in these sites. Thus, formation of protein nanoparticles with as few as four Env trimers was sufficient to significantly alter immunogen trafficking in NHPs and increase delivery to B cell follicles.

## Discussion

Availability of antigen in germinal centers is critical for affinity maturation of the B cell response, but delivery of immunogens to follicles (and subsequent GCs) in a primary immunization is often inefficient for soluble antigens. This is particularly true for soluble HIV Env trimer immunogens, which have been reported in mice to localize to interfollicular macrophages ^20,36^. Efficient immunogen delivery to follicles may be critical because antigens are rapidly degraded *in vivo* ^44^, hence the availability of neutralizing epitopes likely decays over time. Here we tested two different approaches aiming to promote complement-mediated delivery of Env trimer immunogens to follicles, 1) *in vivo* formation of immune complexes with a passively transferred anti-Env monoclonal antibody, and 2) generation of self-assembling protein nanoparticles displaying four copies of stabilized Env trimers. We found that both IC and nanoparticle immunization approaches led to a concentration of antigen ringing the periphery of B cell follicles in the draining lymph nodes of non-human primates. Light sheet imaging of whole cleared lymph nodes revealed that both immunization strategies led to accumulation of the immunogens in follicles throughout the entire lymph node tissue, concentrating in as many as 500 follicles in a single draining lymph node. While both IC and NP immunization have been previously reported to enhance output measures of the humoral immune response to subunit vaccines, our data provides the first demonstration in the closest animal model to humans that these immunization strategies alter immunogen trafficking in a manner that may help drive effective B cell responses.

Extensive progress has been made in the last ten years in the engineering of stable soluble forms of the HIV Env protein that faithfully replicate the antigenic structure of the native viral envelope for potential use in a subunit vaccine ^5–7,9,10,45^. However, it has been shown that glycans of soluble Env trimers are recognized by SIGN-R1-expressing macrophages in lymph nodes, leading to capture of Env immunogens in interfollicular regions of lymph nodes following immunization ^20,36^. Env immunogen accumulation on FDCs has been demonstrated by using novel immunization strategies that deliver antigen delivery over time periods of several weeks through implanted osmotic pumps or repeated injections ^46,47^, but such strategies may not be clinically practical.

Immune complex vaccines have been studied for many years preclinically ^28^ and have also entered clinical trials ^31,32,48^. In the setting of HIV vaccines, complexation with mAbs has been used to alter the accessibility of neutralizing vs. non-neutralizing epitopes or alter conformations of the immunogen, promoting B cell responses against neutralizing epitopes ^28^. Motivated by the recent findings that the base of Env trimers is a highly immunodominant non-neutralizing target on SOSIP immunogens ^46,49,50^, we utilized a mAb isolated from macaques recognizing a base epitope for IC formation, with the goal of masking humoral responses against the base. To allow for the *in vivo* formation of ICs, we preinjected animals intravenously (i.e. passively immunized) with the base-specific RM19R mAb 24 hours prior to antigen administration. While many preclinical studies of IC immunization employ immune complexes formed *ex vivo* prior to injection to conserve antibody ^27,51^, the passive mAb administration used here was selected to both test the impact of monoclonal Ab immune complex formation on immunogen trafficking and model potential effects of boosting animals with pre-existing anti-trimer antibody. Furthermore, we believe this represents an important model for how IC formation affects subunit vaccines, especially for those in which patients likely have pre-existing immunity (e.g. influenza virus and RSV). To our knowledge, this is the first study of the effect of ICs on vaccination in NHPs.

NPs have been reported to exhibit a variety of different distribution patterns *in vivo*. As with immune complexes, innate immunity factors appear to play a significant role in directing the antigen to LN follicles. Link et al. ^52^ showed that Qβ antigens administered to naïve animals only accumulated on the FDC network when they were in a virus-like particle (i.e. a NP) format, not as a soluble dimer. This process was mediated by nIgM and complement. We also recently identified a role for innate immune recognition of particulate HIV antigens in mice ^36^. This pathway involves recognition of the highly-glycosylated residues on Env-based immunogens via mannose-binding lectin (MBL). MBL binding resulted in complement attachment, which directed the NPs to FDCs. Importantly, MBL only bound to NP forms of immunogens, and not soluble subunits, likely due to a requirement for multivalent engagement of MBL with the glycan patches of multiple trimers for stable binding. Due to the success of this approach in mice, we wanted to test if the same principles would hold true in NHPs. Furthermore, to help clarify the degree of antigenic valency that is required for promoting interactions with MBL and directing NP antigen to FDCs, we tested here a new protein NP scaffold that displays only four trimers. This was half the number from the ferritin-scaffold used in the previous mouse study, but the diameters of the two Env-presenting nanoparticles were very similar (~35-40 nm). As a result the spacing between the antigens is almost two times higher in the case of T33_dn2 nanoparticle (~31nm apex-apex distance) versus the ferritin nanoparticle (estimated apex-apex distance for two nearest neighboring trimers ~15-20nm).

We found that both the IC approach and the NP approach were successful in delivering antigen to the FDC network of LN follicles. In both cases, this contrasted to the distribution of soluble Env trimer immunogen in LNs, despite similar or even lower total amounts of antigen accumulation within draining LNs. Concentration of antigen in follicles was observed by multiple methods, including by whole organ clarification and by tissue sectioning and immunofluorescent staining. This was the first time that such a process has been shown in NHPs for Env-presenting NPs, although the pattern of antigen deposition is reminiscent of the localization recently shown for animals receiving sustained antigen delivery via osmotic pumps or repeated injections ^46,47^. The similarity lends support to the hypothesis that ICs from early antigen-specific IgM and IgG were the driving factors for FDC deposition in those “slow delivery” vaccine administration studies.

Antibodies formed in response to slow delivery immunization regimes were found to exhibit enhanced tier 2 HIV virus neutralization and to bind to a more diverse set of epitopes on soluble trimers, which may have been due to the enhanced capture and preservation of the immunogen on FDCs ^46^. We believe that by achieving similar degrees of capture and preservation with bolus administration, it may be possible to evoke similar high-quality antibody responses with ICs or NPs. Additional studies to test for such immune responses are underway.

## Methods

### Env protein production

For immune complex trafficking studies, soluble BG505 SOSIP trimers were employed. The N241 and N289 N-linked glycosylation sites (mutations P240T, S241N, M271I, F288L, T290E and P291S) were introduced into the BG505 SOSIP.v5.2 pPPI4 vector ^53^ using the QuikChange® Lightning Site-Directed Mutagenesis kit (Agilent). The R500A and Q658K mutations were added to the BG505 SOSIP.v5.2.N241.N289 vector. Sequences were verified by Sanger sequencing (Genewiz). BG505 SOSIP.v5.2, BG505 SOSIP.v5.2.N241.N289, and BG505 SOSIP.v5.2.N241.N289.R500A.Q658K were expressed in HEK293F cells (Invitrogen) and purified with PGT145 affinity chromatography followed by size exclusion chromatography (SEC) using a HiLoad® 16/600 Superdex® pg200 (GE Healthcare) as described previously ^53^.

For studies comparing soluble trimer with tetrameric trimer nanoparticles, we employed untagged BG505_SOSIP_MD39 trimers. The protein was expressed in FreeStyle 293-F cells (ThermoFisher Scientific; Cat# R79007) and purified by immuno-affinity purification using 2G12 mAb as previously described ^7^. Briefly, clarified supernatants were passed over 2G12 mAb immobilized on CNBr-Activated Sepharose 4B resin (GE Healthcare; Cat# 17043001). Resin was washed with tris-buffered saline (TBS) and captured trimer was eluted with 3M MgCl2 and immediately buffer-exchanged into fresh TBS. Proteins were concentrated and subjected to size-exclusion chromatography on a HiLoad 26/600 Superdex 200 pg (GE Healthcare; Cat# 28989336). Trimer-containing fractions were pooled, concentrated to 1 mg/mL and flash-frozen on liquid nitrogen in 100 µL aliquots and stored at −80C until further use. All steps were performed under low-endotoxin conditions and the final preparation was confirmed to be <2.5 EU/mg using an Endosafe-PTS system (Charles River; Cat# PTS100).

### RM19R IgG and Fab production

The sequences encoding rhesus macaque IgG1 heavy chain of RM19R and the rhesus macaque kappa chain of RM19R were codon optimized, synthesized, and separately cloned into the pcDNA3.4 vector using the GeneArt service from Invitrogen. The RM19R IgG was produced by transient transfection of Expi293 cells and purified using Protein A affinity chromatograph by Invitrogen. The RM19R Fab heavy chain plasmid was made introducing two stop codons following residue D234 in the RM19R IgG1 heavy chain vector using the QuikChange® Lightning Site-Directed Mutagenesis kit (Agilent). The RM19R Fab was expressed in HEK293F cells (Invitrogen) by co-transfecting with the Fab heavy and kappa chain plasmids (1:1 ratio) using PEImax. Transfection supernatant was harvested after 6 days and passed through a 0.45 µm filter. The RM19R Fab was purified using CaptureSelect™ CH1-XL (ThermoFisher) affinity chromatography.

### BG505 SOSIP-T33_dn2 nanoparticle component production

BG505-SOSIP was engineered with stabilizing mutations from SOSIP.v5.2 ^53^ (A501C, T605C, I559P, E64K, A73C, A316W, A561C) and MD39 ^9^ (M271I, A319Y, R585H, L568D, V570H, R304V, F519S) and glycan knock-ins at positions N241 (mutations: P240T, S241N) and N289 (mutations: F288L, T290E, P291S) was fused to component A of the T33_dn2 nanoparticle and subcloned into a pPPI4 vector as described previously ^38^. The vector with antigen-bearing component was transfected into FreeStyle 293F cells using polyethylenimine (Polysciences, Inc) as described previously ^7^. Six days post transfection the cells were removed by centrifugation (7000 RPM for 1 hour at 4 °C) and the supernatant was cleared by vacuum filtration (0.45 µm filtration units, Millipore Sigma). BG505 SOSIP-T33_dn2A component was purified from the cleared supernatant using Sepharose 4B resin (GE Healthcare Life Sciences) carrying PGT145 IgG. The resin was washed with buffer containing 25 mM Tris-HCl + 500 mM NaCl (pH 7.4) and the protein was eluted using buffer containing 3 M MgCl_2_ + 250 mM l-arginine (pH 7.2). Eluate was collected into an equal volume of the size exclusion chromatography (SEC) buffer (25 mM Tris + 500 mM NaCl + 250 mM l-arginine + 5 % glycerol, pH 7.4). The sample was concentrated and buffer exchanged to the SEC buffer using Amicon ultrafiltration units with 100 kDa cutoff (Millipore Sigma). A HiLoad 16/600 Superdex S200 pg column was used for the gel filtration step. The protein was concentrated and stored in SEC buffer at 4 °C until nanoparticle assembly. The T33_dn2B component of the nanoparticle was expressed in E coli. BL21-DE3 cells (NEB) were transformed with pET28b vector carrying T33_dn2B gene with a C-terminal His-tag. Following inoculation the cells were incubated in self-inducible media ^37^ and shaken at 220 RPM at 16 °C for ~18 hours. BL21 cells were spun down (3000 RPM, 30 min, 4 °C) and resuspended in Tris-buffered saline (TBS, Alfa Aesar) containing cOmplete™ protease inhibitor cocktail (Sigma Millipore). The cells were lysed using sonication and pressurized disruption. Cell lysate was cleared by centrifugation at 12000 RPM for 1 hour at 4 °C, filtered (0.45 µm filtration units, Millipore Sigma) and loaded onto a cOmplete™ His-Tag Purification Resin gravity column (Sigma Millipore). Resin was first washed with detergent-containing buffer (25 mM Tris + 500 mM NaCl + 0.5 % N-Dodecyl-β-D-maltoside, pH 7.2) and then with low imidazole buffer (25 mM Tris + 500 NaCl + 20 mM Imidazole, pH 7.2). Detergent buffer wash helped remove endotoxin from the sample. Sample was eluted using high imidazole buffer (25 mM Tris + 500 NaCl + 500 mM Imidazole, pH 7.2), concentrated and buffer-exchanged to the same SEC buffer as described above using Amicon ultrafiltration units with 10 kDa cutoff (Millipore Sigma). Finally, T33_dn2B was SEC purified using HiLoad 16/600 Superdex S200 pg column.

### Nanoparticle assembly, purification and labeling

Nanoparticle components (BG505 SOSIP-T33_dn2A and T33_dn2B) were concentrated to ~1 mg/ml and equimolar amounts were combined and incubated for 24 hours at 4 °C, for nanoparticle assembly. Assembled nanoparticles were purified from the unassembled components using Sephacryl S-500 HR column using DPBS (Thermo Fisher Scientific) as the running buffer. ToxinSensorTM Single Test Kit (GenScript) was applied to verify that the endotoxin levels of the labeled nanoparticle were below 50 EU/kg per dose.

### Fluorescent labeling

RM19R IgG was labeled with the Alexa Fluor™ 647 Protein Labeling Kit (Thermo Fisher) to a degree of labeling (DoL) of 7.1 fluorophores per Ab. The BG505 SOSIP.v5.2.N241.N289 trimer was labeled with the Alexa Fluor™ 568 Protein Labeling Kit (Thermo Fisher) to a DoL of 8.3 fluorophores per trimer. The MD39 SOSIP trimer was labeled with the Alexa Fluor™ 647 Protein Labeling Kit (Thermo Fisher) to a degree of labeling (DoL) of 4.1 fluorophores per trimer. For the BG505 SOSIP-T33_dn2 nanoparticle, 2 mg of concentrated nanoparticles were labeled using an Alexa Fluor™ 647 Protein Labeling Kit (Thermo Fisher). The final degree of labeling (DoL) was 42.1 fluorophores per nanoparticle; 10.5 per trimer.

### Bio-Layer Interferometry (BLI)

An Octet RED instrument (ForteBio) was used to determine the kinetic parameters of the RM19R/BG505 SOSIP interaction by Biolayer Interferometry. The RM19R Fab was loaded onto anti-human Fab-CH1 (FAB2G) biosensors (ForteBio) at a concentration of 10 μg/mL in kinetics buffer (PBS, pH 7.4, 0.01% (w/v) BSA, and 0.002% (v/v) Tween 20) until response of 1 nanometer shift was reached. The loaded biosensors were dipped into kinetics buffer for 1 min to acquire a baseline and then moved to wells containing a series of 2-fold dilutions of BG505 SOSIP.v5.2 in kinetics buffer, starting at a 125 nM. The trimers were allowed to associate for 180 secs before the biosensor were move back to the wells containing kinetics buffer where the baseline was acquired. Dissociation of the trimers from the Fab-loaded biosensors was recorded for 300 secs. Kinetic parameters were calculated using the Octet System Data Analysis v9.0 (ForteBio). To assess how the R500A and Q658K mutations impact binding of RM19R the IgG was loaded onto anti-human IgG Fc capture (AHC) biosensors (ForteBio) at a concentration of 5 μg/mL in kinetics buffer until response of 1 nanometer shift was reached. The loaded biosensors were dipped into kinetics buffer for 1 min to acquire a baseline and then moved to wells containing either BG505 SOSIP.v5.2 or BG505 SOSIP.v5.2.N241.N289.R500A.Q658K, both at a concentration of 1000 nM in kinetics buffer. The trimers were allowed to associate for 180 secs before the biosensors were moved back to the wells containing kinetics buffer where the baseline was acquired. Dissociation of the trimers from the IgG-loaded biosensors was recorded for 180 secs.

### Negative stain electron microscopy

Negative stain electron microscopy (NS-EM) experiments were performed as described previously ^54,55^. Fusion components and assembled nanoparticle samples were diluted to 20-50 µg/ml and loaded onto the carbon-coated 400-mesh Cu grid that had previously been glow-discharged at 15 mA for 25 s. Grids were negatively stained with 2 % (w/v) uranyl-formate for 60 s. Data collection was performed on either an FEI Tecnai T12 microscope (2.05 Å/pixel; 52,000× magnification) or FEI Talos microscope (1.98 Å/pixel; 72,000× magnification). The electron dose was set to 25 e-/Å^2^ and images were collected with a defocus value of −1.50 or −2.00 µm. The micrographs were recorded on a Tietz 4k x 4k TemCam-F416 CMOS or a FEI Ceta 16M camera using a Leginon automated imaging interface ^56^. Data processing was performed in Appion data processing suite. With nanoparticle samples, approximately 500 – 1000 particles were manually picked from the micrographs and 2D-classified using the Iterative MSA/MRA algorithm. With trimer samples, 10,000 – 40,000 particles were auto-picked and 2D-classified using the Iterative MSA/MRA algorithm. For 3D classification and refinement, we continued processing in Relion/3.0b3 ^57^.

### BG505 SOSIP/RM19R Fab complex

700 μg of the BG505 SOSIP.v5.2.N241.N289 trimer was mixed with approximately 1.4 mg RM19R Fab and incubated at RT overnight. The complex was SEC purified using a Superose™ 6 Increase 10/300 GL (GE Healthcare) column in TBS. Fractions containing the complex were concentrated to 13 mg/mL using a 100 kDa Amicon® spin concentrator (Millipore).

### Cryo-EM grid preparation

A Vitrobot mark IV (Thermo Fisher Scientific) was used for grid preparation for both the BG505 SOSIP/RM19R complex and BG505 SOSIP-T33_dn2 nanoparticle samples. Temperature was set to 10 °C, humidity at 100% with a 4-7 s blotting time, blotting force of 0 and wait time of 10 s. For the RM19R sample, 3 μL of the BG505 SOSIP/RM19R complex at 13 mg/mL was mixed with 1 μL of a n-Dodecyl-β-D-Maltopyranoside (DDM) solution to a final DDM concentration of 0.06 mM and applied to a C-Flat grid (CF-2/2-4C, Protochips, Inc.), which had been plasma-cleaned for 5 seconds using a mixture of Ar/O_2_ (Gatan Solarus 950 Plasma system). For the BG505 SOSIP-T33_dn2 nanoparticle sample, lauryl maltose neopentyl glycol (LMNG) at a final concentration of 0.005 mM was added to the nanoparticle sample (4.0 mg/ml) and 3 µl was immediately loaded onto plasma cleaned Quantifoil R 2/1 holey carbon copper gri (Cu, 400-mesh, Quantifoil Micro Tools GmbH). Blotted grids were plunge-frozen into nitrogen-cooled liquid ethane.

### Cryo-EM data collection and processing

Samples were imaged on either FEI Titan Krios electron microscope (ThermoFisher) operating at 300 keV (RM19R dataset) or a FEI Talos Arctica electron microscope (ThermoFisher) operating at 200 keV (T33_dn2 nanoparticle datasets). Both microscopes were equipped with Gatan K2 Summit direct electron directors operating in counting mode. Automated data collection was performed using the Leginon software suite ^56^. Micrograph movie frames were aligned and dose-weighted using MotionCor2 ^58^, and CTF models were determined using Gctf ^59^. For the RM19R dataset, dose-weighted micrographs were assessed using EMHP ^60^ and particles were picked using DoG Picker ^61^. Particle extraction, 2D classification, and 3D refinement were conducted using Relion v2.1 ^62^. For the T33_dn2 nanoparticle, 3 datasets were collected and initial data processing was performed in cryoSPARC v2.5.0 ^63^. Particles were picked using cryoSPARC template picker, extracted and 2D classified. 89,863 particles belonging to nanoparticle classes were then transferred to Relion/3.0 ^57^ for further processing. A reference model was generated using Ab-Initio Reconstruction in cryoSPARC. Several iterative 3D refinement and classification steps in Relion were applied to identify a subpopulation of 35,521 particles that went into the final 3D reconstructions. Tetrahedral symmetry was applied for all 3D steps. Soft solvent mask around the nanoparticle core was introduced during the final 3D classification, refinement and post-processing steps. Final resolution of the NP core was 4.6 Å after post-processing. Trimeric BG505 SOSIP antigens were connected to the nanoparticle core via a flexible linker and they appeared disordered in the final map of the nanoparticle. In order to obtain high-resolution information on the BG505 SOSIP trimers we applied localized reconstruction v1.2.0 ^64^. First, marker files were generated in UCSF Chimera ^65^ to define the subparticle vectors. Then the trimer subparticles were extracted from aligned nanoparticles from which the signal corresponding to the nanoparticle core has been subtracted. Each nanoparticle displayed four trimers so the total number of extracted subparticles was 142,084 (4 x 35,521). Trimer subparticles were then 2D and 3D classified in Relion. A subset of 52,939 subparticles was subjected to 3D auto refinement with C3 symmetry. Soft solvent mask was applied during refinement and postprocessing steps. Final resolution of the presented BG505 SOSIP antigen map is 4.5 Å. A graphical summary of the data processing approach and relevant statistics are displayed in Supplementary Figure 1.A. Data collection and processing parameters are reported in **Supplementary Table I**.

### Model building and refinement

A model of the Fv region of RM19R was generated using the Rosetta antibody protocol available on the ROSIE server ^66,67^. An initial molecular model of the BG505 SOSIP trimer/RM19R Fab complexes was built by docking the Env portion of PDB: 5ACO ^68^ into the EM density maps along with the RM19R Fv model generated in Rosetta using UCSF Chimera ^65^. The appropriate mutations were introduced into the Env sequence to match the BG505 SOSIP.v5.2.N241.N289 sequence and N-linked glycans were added using Coot ^69^. Initial models for T33_dn2 nanoparticle core and the BG505 SOSIP trimer were relaxed into the post-processed, B-factor sharpened maps following 3D refinement. For all models, iterative rounds of Rosetta relaxed refinement ^70,71^ and manual refinement in Coot ^69^ were used to generate the final models. Appropriate symmetry (tetrahedral for NP and C3 for BG505 SOSIP trimer and BG505 SOSIP/RM19R Fab complex) was applied during the automated refinement steps. EMRinger and MolProbity scores were used for assessment of refined models ^72,73^. Glycan structures were validated by CARP ^74^, pdb-care ^75^, and Privateer ^76^. Final model statistics are summarized in **Supplementary Table II**.

### Immunizations

#### Animals

Indian rhesus macaques (*Macaca mulatta*) (RM) were housed at the Yerkes National Primate Research Center and maintained in accordance with NIH guidelines. This study was approved by the Emory University Institutional Animal Care and Use Committee (IACUC). All animals were treated with anesthesia and analgesics for procedures as per veterinarian recommendations and IACUC approved protocol.

#### Adjuvant

The adjuvant used for all immunizations was an ISCOM-like nanoparticle comprised of self-assembled cholesterol, phospholipid, and Quillaja saponin prepared as previously described ^77^.

#### Immune complex study

Pairs of RM were injected i.v. with 1:1 mixtures of fluorescently-labeled and unlabeled RM19R mAb at 2.0 mg/kg or 0.2 mg/kg, and 24 hr later, all animals received s.c. injections in each thigh of 50 µg BG505 SOSIP trimer immunogen combined with 187.5 µg saponin adjuvant; a control group received SOSIP trimer and adjuvant in the absence of RM19R passive transfer. Three days after antigen injection, animals were sacrificed and all LNs in selected locations were collected for analysis.

#### Nanoparticle immunogen study

Groups of three RM were immunized on day 0 with 50 µg BG505 SOSIP MD39 trimer ^9^ or 71 µg BG505 SOSIP-T33_dn2 nanoparticle (to achieve an equimolar dose on a per-trimer basis) conjugated to Alexa Fluor 647 s.c. in both left and right inner mid thighs combined with 187.5 µg of saponin adjuvant. Animals were sacrificed two days or seven days after immunization and all LNs in selected locations were collected for analysis.

### Ex vivo tissue fixation

All NHP LNs were harvested and immediately placed in PLP buffer (pH 7.4 50 mM PBS + 100 mM lysine, 1% paraformaldehyde, 2 mg/mL sodium periodate) for fixation. After 4-5 days at 4°C, the tissues were washed and stored in PBS with 0.05% sodium azide at 4°C until taken for imaging.

### Whole organ fluorescence measurement

Total antigen signal within LNs from the antigen tracking study was measured in two ways. First, by IVIS (In Vivo Imaging System, PerkinElmer) fluorescence imaging of whole tissues within clear plastic 24-well plates, using excitation at 570 nm and emission at 620 nm (for trimer-AF568) and excitation at 640 nm and emission at 680 nm (for RM19R-AF647). Alternatively, fluorescence was measured by placing the tissues directly on the glass scanning surface of a Typhoon FLA 9500 biomolecular imager (GE Healthcare Life Sciences) and using either a 535 nm excitation laser with a >575 nm long-pass filter, or a 635 nm excitation laser and a ≥665 nm long-pass filter. The integrated signal density corresponding to labeled antigen or antibody in each LN was calculated using LivingImage (PerkinElmer) or ImageJ software and plotted using GraphPad Prism 8. For comparing trimer to nanoparticle in Fig. 5B, the difference in degree of labeling between soluble trimer and the nanoparticle was used to normalize the signal on a per-trimer basis.

### Whole organ clarification

Selected LNs were clarified via a combination/modification of the iDISCO ^40^ and CUBIC ^41^ organ-clearing methods. The LNs were first delipidated based on the iDISCO methanol incubation protocol: Tissues were washed in water for 1 hour, then 20% methanol in water for 2 hours. A series of stepwise increases in methanol percentage (40%, 60%, 100%, 100%) followed, each step for 2 hours. The LNs were then placed into 2:1 MeOH:DCM overnight, and the next day were rehydrated with the following series of methanol solutions for 2 hours each: 100%, 100%, 80%, 60%, 40%, 20%, 0%, 0%. Next, the LNs were placed into 10-20 mL of a 1:1 mixture of CUBIC-R index-matching solution (45 w% antipyrine and 30 w% nicotinamide in water) for 1 day, followed by at least 20 mL of undiluted CUBIC-R for 2 days or as long as needed for adequate clarification. Larger organs were moved into a fresh 20 mL of CUBIC-R solution to ensure that the refractive index of the solution would not be significantly lowered by residual water in the tissue.

### Light sheet microscopy

Clarified LNs were imaged in CUBIC-R solution using a LaVision Ultramicroscope II Light Sheet Microscope at 1.25x optical zoom. The Alexa Fluor 647-labeled antigen or antibody was imaged using the 640 nm laser and the Alexa Fluor 568-labeled antigen was imaged using the 561 nm laser at 100 ms exposure time on an Andor Neo camera. Snapshots and movies were generated using the 3D viewer in the FIJI package of ImageJ. Follicles containing fluorescently-labeled antigen were counted manually using the maximum projection of each LN.

### Immunofluorescence staining

Selected LNs were embedded in 3% low melting temperature agarose (Sigma-Aldrich), and then sliced into 350 µm-thick sections using a vibratome. The slices were blocked and permeabilized overnight in PBS with 5% mouse serum, 5% rat serum, and 0.2% Triton-X-100, followed by staining for 3 days at 37 °C with 1:100 dilutions of BV421-labeled mouse anti-human CD35 clone E11 (BD Biosciences) and Alexa Fluor 488-labeled mouse anti-Ki67 clone B56 (BD Biosciences) in the same buffer as the blocking/permeabilization step. Stained slices were then washed for 3 days at room temp with PBS containing 0.2% Tween-20, and then mounted onto glass slides with Prolong Glass antifade mountant (Thermo Fisher Scientific).

### Confocal microscopy

Imaging was performed on either one of a Leica SP8 or an Olympus FV1200 laser scanning confocal microscope using 10x air or 20x water immersion objectives. Images were analyzed using ImageJ.

### Binding experiments with Mannose-Binding Lectin (MBL)

Truncated human MBL-2 was purchased from MyBioSource (Catalog # MBS2086086). 5 µg of MBL-2 was incubated with 5 µg of BG505 SOSIP or BG505 SOSIP-T33_dn2 nanoparticle, for 4 hours at 37 °C in TBS (Alfa Aesar) buffer containing 2 mM CaCl_2_. Following the incubation, the samples were diluted to 40 µg/ml, loaded onto carbon-coated copper EM grids, negatively stained with uranyl-formate and imaged as described in the negative-stain EM methods section. Imaging was also performed on grids containing free, non-complexed MBL-2, BG505 SOSIP and BG505 SOSIP-T33_dn2 NP for comparison. For size-exclusion chromatography experiments, 30 µg of MBL-2 was mixed with 15 µg of BG505-SOSIP or BG505 SOSIP-T33_dn2 nanoparticle samples and incubated for 4 hours at 37 °C in TBS + 2 mM CaCl_2_. For comparison, equivalent amounts of free MBL-2, BG505-SOSIP and BG505 SOSIP-T33_dn2 nanoparticle were incubated under identical conditions. For size-exclusion chromatography step (SEC), we used Superose 6 Increase 10/300 column, running in TBS + 2 mM CaCl_2_. SEC traces of the assembly and control reactions are shown in **Supplementary Fig. 4B** and C.

### Data Availability

Cryo-EM reconstructions have been deposited in the Electron Microscopy Data Bank (EMDB: EMD-21227, EMD-21230, and EMD-21231), and in the Protein Data Bank (PDB: 6VKN, 6VL5, and 6VL6).

## Acknowledgments

We thank Jean-Christophe Ducom, Hannah L. Turner, Bill Anderson, and Charles Bowman for assistance with computational resources, microscope management, and data collection. This work was supported in part by the NIH (awards UM1AI100663 and UM1AI144462 to D.J.I., S.C., W.R.S., G.S. and A.B.W., AI125068 to D.J.I., S.C., and G.S., and P01AI048240 to D.J.I.), the Bill & Melinda Gates Foundation (OPP1156262 to D.B. and N.P.K.), and the Ragon Institute for MGH, MIT, and Harvard. C.A.C. is supported by the NIH F31 Ruth L. Kirschstein Predoctoral Award Al131873 and by the Achievement Rewards for College Scientists Foundation. The content is solely the responsibility of the authors and does not necessarily represent the official views of the National Institutes of Health. D.B and D.J.I. are investigators of the Howard Hughes Medical Institute.

## Author Contributions

A.B.W., S.C., D.J.I., J.T.M., K.C., A.A., and C.A.C. conceived the study plan. J.T.M. performed the mouse experiments. J.T.M. and B.J.C. fluorophore-labeled BG505-SOSIP and performed all fluorescent imaging experiments. C.A.C. produced and characterized RM19R, fluorophore-labeled RM19R, and performed ELISAs. A.A. and J.C. produced, characterized, and fluorophore-labeled the T33_dn2 nanoparticle, and carried out MBL binding studies. G.U., N.P.K., and D.B. designed the T33_dn2 nanoparticle. T.T. produced the ISCOM-like adjuvant. D.G.C., C.A.E., E.H.G, Y.C., F.V., and G.S. performed the NHP immunizations and tissue collection. T.S, S.M., and W.R.S. provided the BG505 SOSIP trimer used in the nanoparticle comparison study. J.T.M., C.A.C., A.A., A.B.W., and D.J.I. wrote the manuscript. A.B.W., G.S., and D.J.I. supervised the study.

