## Supplementary Information for "Targeting HIV Env immunogens to B cell follicles in non-human primates through immune complex or protein nanoparticle formulations"

**Supplementary Table I.** Cryo-EM data collection information

| | BG505 SOSIP + RM19R Fab | BG505 SOSIP-T33_dn2 nanoparticle | |
|---|---|---|---|
| **Microscope** | Titan Krios | Talos Arctica | |
| **Voltage (kV)** | 300 | 200 | |
| **Detector** | Gatan K2 | Gatan K2 Summit | |
| **Recording mode** | Counting | Counting | |
| **Magnification** | 29,000 X | 36,000 X | |
| **Movie micrograph pixel size** | 1.03 | 1.15 | |
| **Dose rate ($e^-$/Å$^2$/s)** | 5.69 | 4.44 | |
| **No. of frames per movie** | 50 | 45 | |
| **Frame exposure time (ms)** | 250 | 250 | |
| **Movie micrograph exposure time** | 12.50 | 11.25 | |
| **Total dose ($e^-$/Å$^2$)** | 67.0 | 50.0 | |
| **Under focus range (μm)** | 0.9 – 2.2 | 0.8 – 2.0 | |
| **Number of movie micrographs** | 1247 | 2748 | |
| **Resolution (Å)** | 3.71 | 4.60* | 4.46[#] |
| **Number of particles** | 191556 | 35521* | 52939[#] |
| **EMDB** | EMD-21227 | EMD-21231* | EMD-21230[#] |

* Values/entries refer to the T33_dn2 nanoparticle core subparticle.
[#] Values/entries refer to the BG505 SOSIP antigen subparticle.

**Supplementary Table II.** Model refinement statistics

| | BG505 SOSIP + RM19R | T33_dn2 Nanoparticle core | BG505 SOSIP Trimer |
|---|---|---|---|
| **PDB ID** | 6VKN | 6VL6 | 6VL5 |
| **Residues** | 2496 | 2640 | 1827 |
| **Amino-acids** | 2394 | 2640 | 1719 |
| **Carbohydrates** | 102 | 0 | 108 |
| **RMSD Bonds** | 0.018 | 0.019 | 0.018 |
| **RMSD Angles** | 1.690 | 1.457 | 1.712 |
| **Ramachandran** | | | |
| **Favored (%)** | 95.65 | 99.07 | 98.75 |
| **Allowed (%)** | 3.58 | 0.93 | 1.25 |
| **Outliers (%)** | 0.77 | 0.00 | 0.00 |
| **Rotamer outliers (%)** | 0.85 | 0.00 | 0.20 |
| **Clash score** | 1.05 | 0.28 | 0.61 |
| **Molprobity score** | 1.11 | 0.61 | 0.70 |
| **EMRinger score** | 3.33 | 0.21 | 1.55 |

**Supplementary Figure 1:**

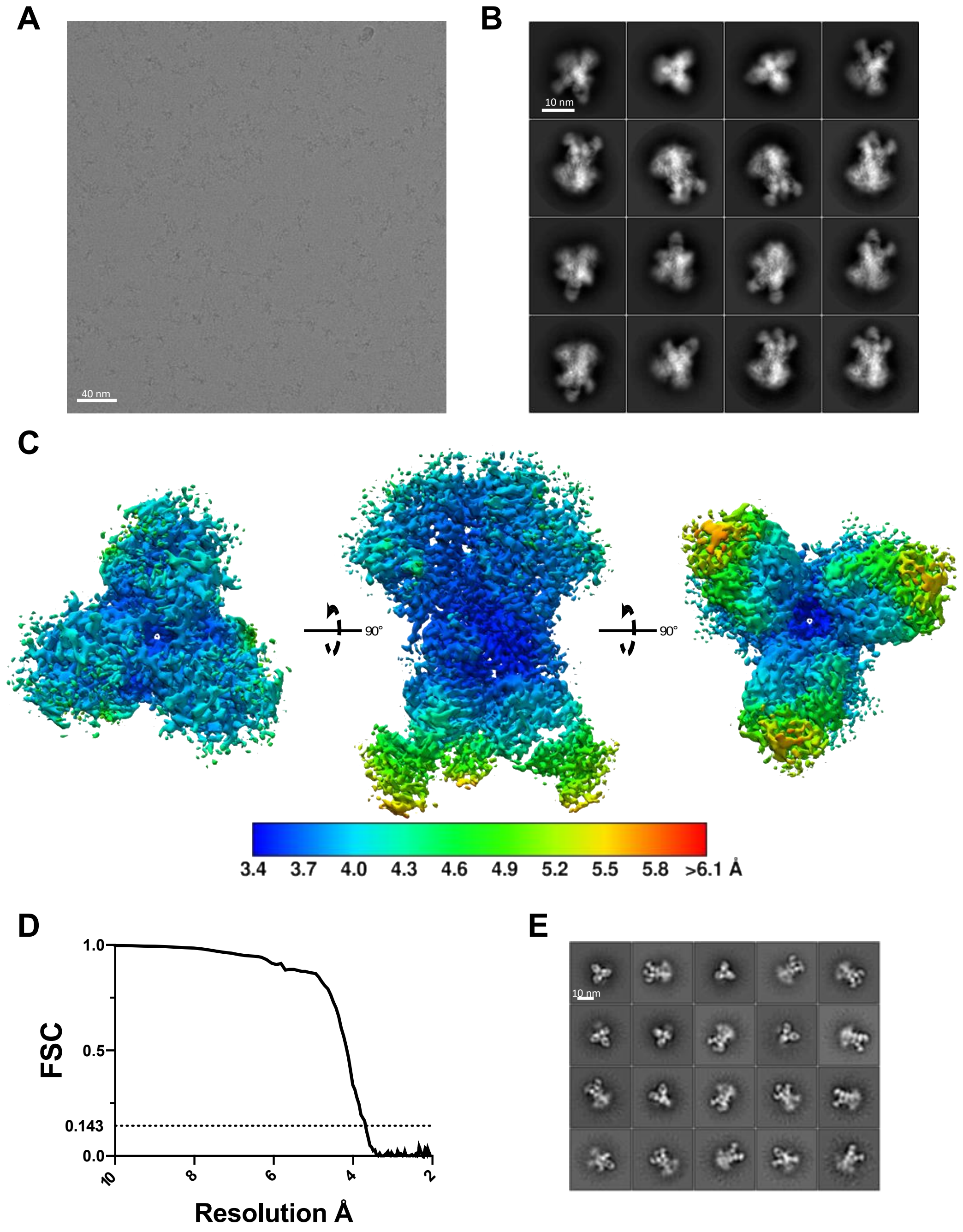


**Supplementary Figure 1.**

(A) Representative cryoEM micrograph of BG505 SOSIP.v5.2.N241.N289 trimer/RM19R Fab complex.

(B) 2D class averages of BG505 SOSIP.v5.2.N241.N289 trimer/RM19R Fab complex.

(C) Local resolution maps for BG505 SOSIP.v5.2.N241.N289 trimer/RM19R Fab complex.

(D) Gold-standard Fourier shell correlation (FSC) curves for BG505 SOSIP.v5.2.N241.N289 trimer/RM19R Fab complex showing global resolution calculated at FSC = 0.143.

(E) NS-EM 2D class averages of BG505 SOSIP.v5.2 following incubation with RM19R Fab at RT for >24 hrs.

**Supplementary Figure 2:**

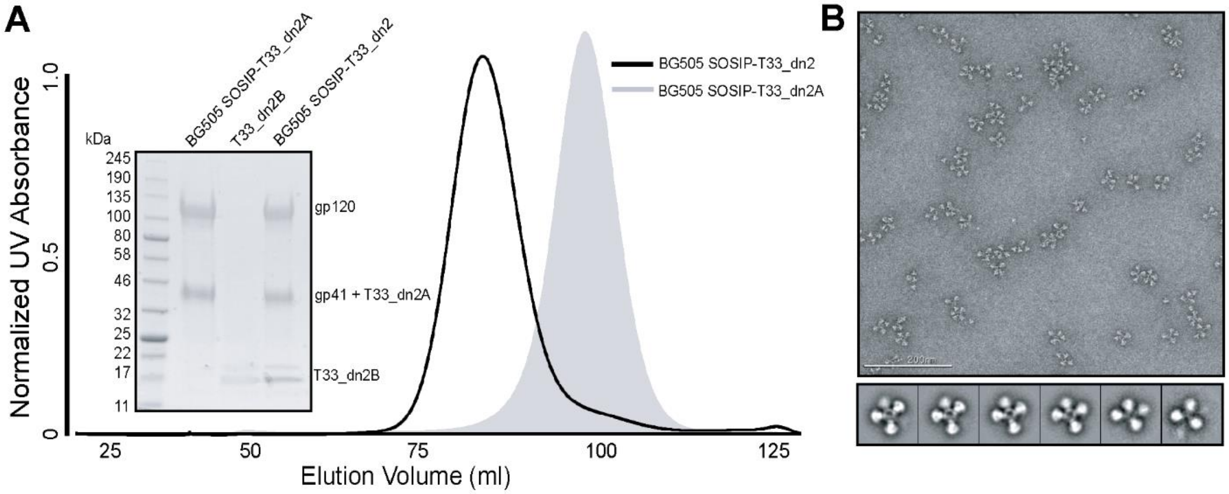


**Supplementary Figure 2.**

(A). Overlay of SEC chromatograms corresponding to BG505 SOSIP-T33_dn2A and assembled BG505 SOSIP-T33_dn2 nanoparticle and an SDS PAGE gel of the purified nanoparticle sample (B). Representative negative stain EM micrograph and 2D class averages of the assembled nanoparticle

### Supplementary Figure 3:

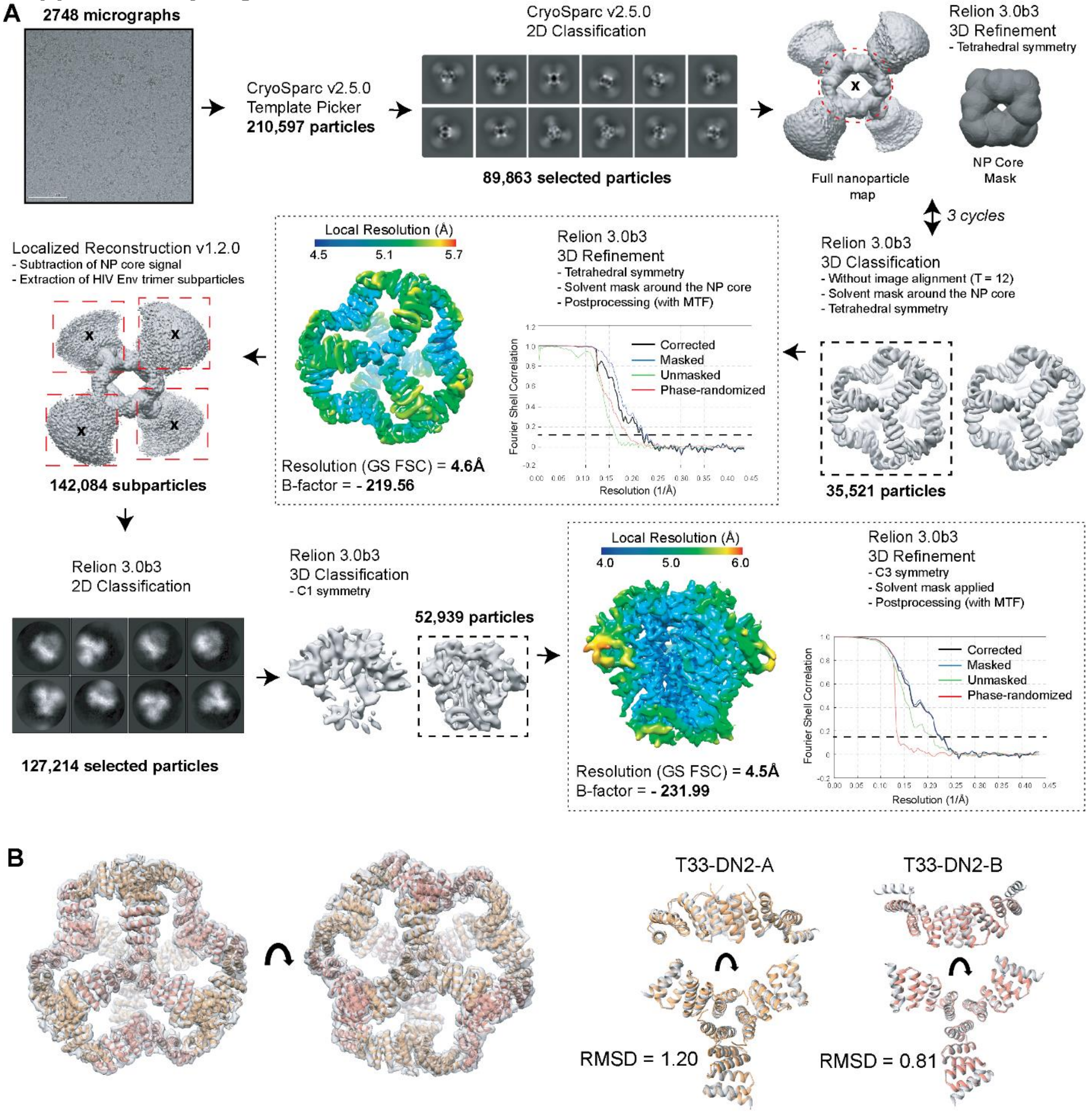


**Supplementary Figure 3.**

Cryo-EM data processing workflow for BG505 SOSIP-T33_dn2 nanoparticle dataset with relevant statistics (A). Fit of refined nanoparticle model to reconstructed density (left) and overlay of refined and Rosetta_design-predicted model of T33_dn2A and T33_dn2B (right) (B).

### Supplementary Figure 4:

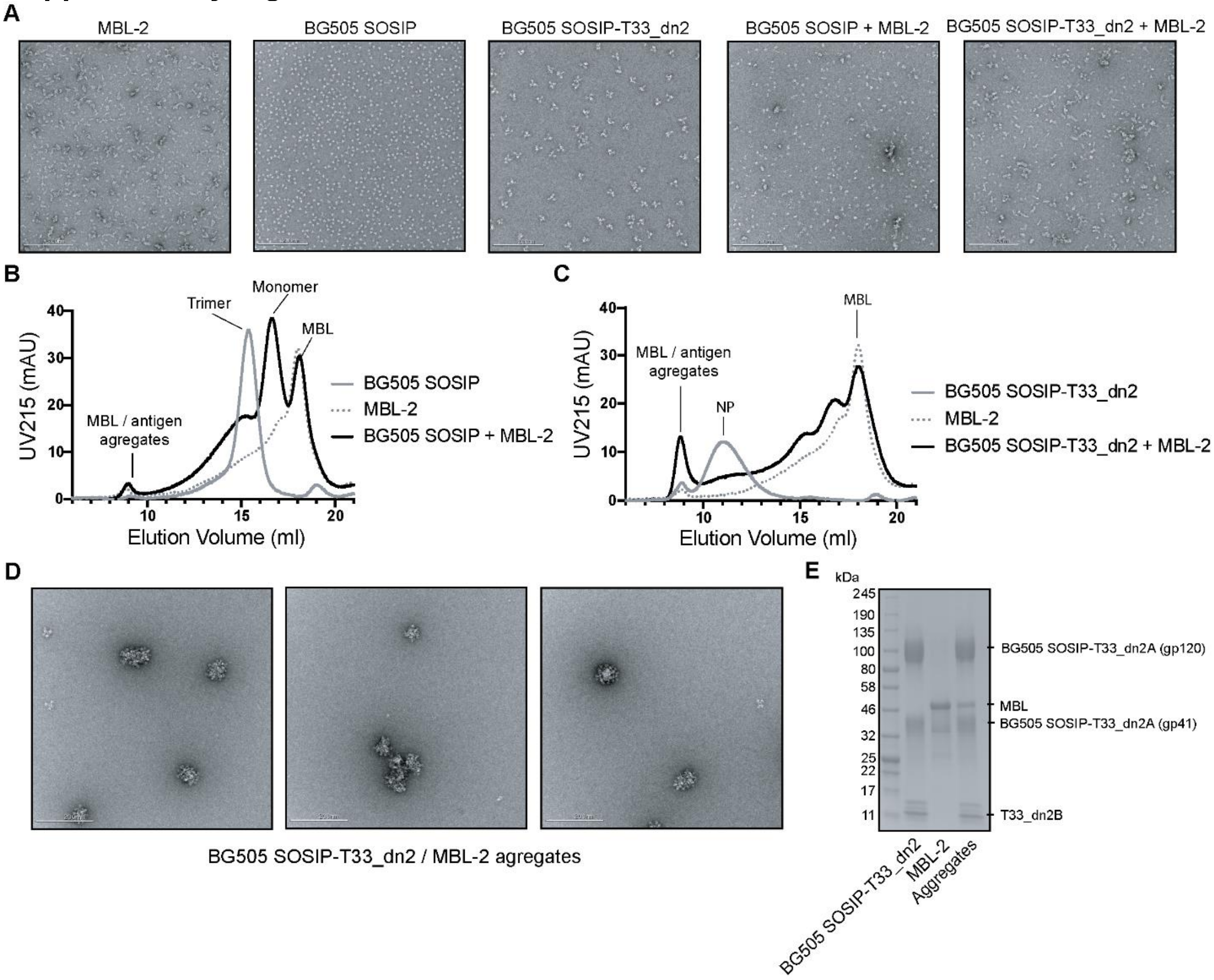


**Supplementary Figure 4. MBL binding experiments.**

(A) Human MBL-2 was incubated with free BG505 SOSIP and BG505 SOSIP-T33_dn2 nanoparticles for 4 hours at 37 °C. Negative-stain EM micrographs of the two samples are shown on the right. For comparison, EM micrographs of free MBL-2, BG505-SOSIP trimer and BG505 SOSIP-T33_dn2 nanoparticle are shown on the left. (B) SEC traces of free MBL-2, BG505 SOSIP and their combination following a 37 °C incubation. (C) SEC traces of free MBL-2, BG505 SOSIP-T33_dn2 nanoparticle and the combined sample following a 37 °C incubation. (D) SEC fractions corresponding to high-molecular weight aggregates in the MBL-2 + BG505 SOSIP-T33_dn2 sample were pooled, concentrated and imaged using NS-EM. Representative micrographs are shown. (E) SDS PAGE gel of free BG505 SOSIP-T33_dn2 nanoparticle, MBL-2 and the BG505 SOSIP-T33_dn2/MBL-2 aggregates purified by SEC.

**Supplementary Video 1. 3D perspective and 360° rotation of cleared LNs from RM19R immune complex study.**
Samples & data are the same as in **Fig. 3D**, displayed in three dimensions with rotation. Scale bars are 1 mm.

**Supplementary Video 2. Close-up on follicles from cleared LNs from RM19R immune complex study, showing bowl-shaped morphology of follicular accumulation.**
Samples are the same as in **Fig. 3D**, imaged at higher magnification (12.5x). Scale bars are 100 µm. Apparent changes in brightness during rotation are due to software interpolation in the z-plane (optical sections are spaced every 5 µm, resolution in x & y is 0.52 µm / pixel).

**Supplementary Video 1. 3D perspective and 360° rotation of cleared LNs from trimer vs. nanoparticle comparison study.**
Samples & data are the same as in **Fig. 5C**, with rotation. Scale bar is 5 mm.
